# Extracellular Matrix Stiffness Promotes Vascular Smooth Muscle Cell Calcification By Reducing The Levels Of Nuclear Actin Monomers

**DOI:** 10.1101/2023.06.18.545506

**Authors:** MC McNeill, F Li Mow Chee, R Ebrahimighaei, GB Sala-Newby, AC Newby, T Hathway, AS Annaiah, S Joseph, M Carrabba, M Bond

**Affiliations:** Department of Translational Health Sciences, Bristol Medical School, Bristol, BS2 8HW; Bristol Heart Institute, University Hospital, Bristol NHS Foundation Trust, Bristol, BS2 8HW

**Keywords:** Calcification, Vascular smooth muscle cell, stiffening, TEAD, RUNX2, extracellular matrix

## Abstract

1.0

**Background:** Vascular calcification (VC) is a prevalent independent risk factor for adverse cardiovascular events and is associated with diabetes, hypertension, chronic kidney disease, and atherosclerosis. However, the mechanisms regulating the osteogenic differentiation of vascular smooth muscle cells (VSMC) are not fully understood.

**Methods:** Using hydrogels of tuneable stiffness and lysyl oxidase-mediated stiffening of human saphenous vein ex vivo,we investigated the role of extracellular matrix (ECM) stiffness in the regulation of VSMC calcification

**Results:** We demonstrate that increased ECM stiffness enhances VSMC osteogenic differentiation and VSMC calcification. We show that the effects of ECM stiffness are mediated via a reduction in the level of actin monomer within the nucleus. We show that in cells interacting with soft ECM, elevated levels of nuclear actin monomer repress osteogenic differentiation and calcification by repressing YAP-mediated activation of both TEA Domain transcription factor (TEAD) and RUNX Family Transcription factor 2 (RUNX2).

**Conclusion:** This work highlights for the first time the role of nuclear actin in mediating ECM stiffness-dependent VSMC calcification and the dual role of YAP-TEAD and YAP-RUNX2 transcriptional complexes.

**GRAPHICAL ABSTRACT:** 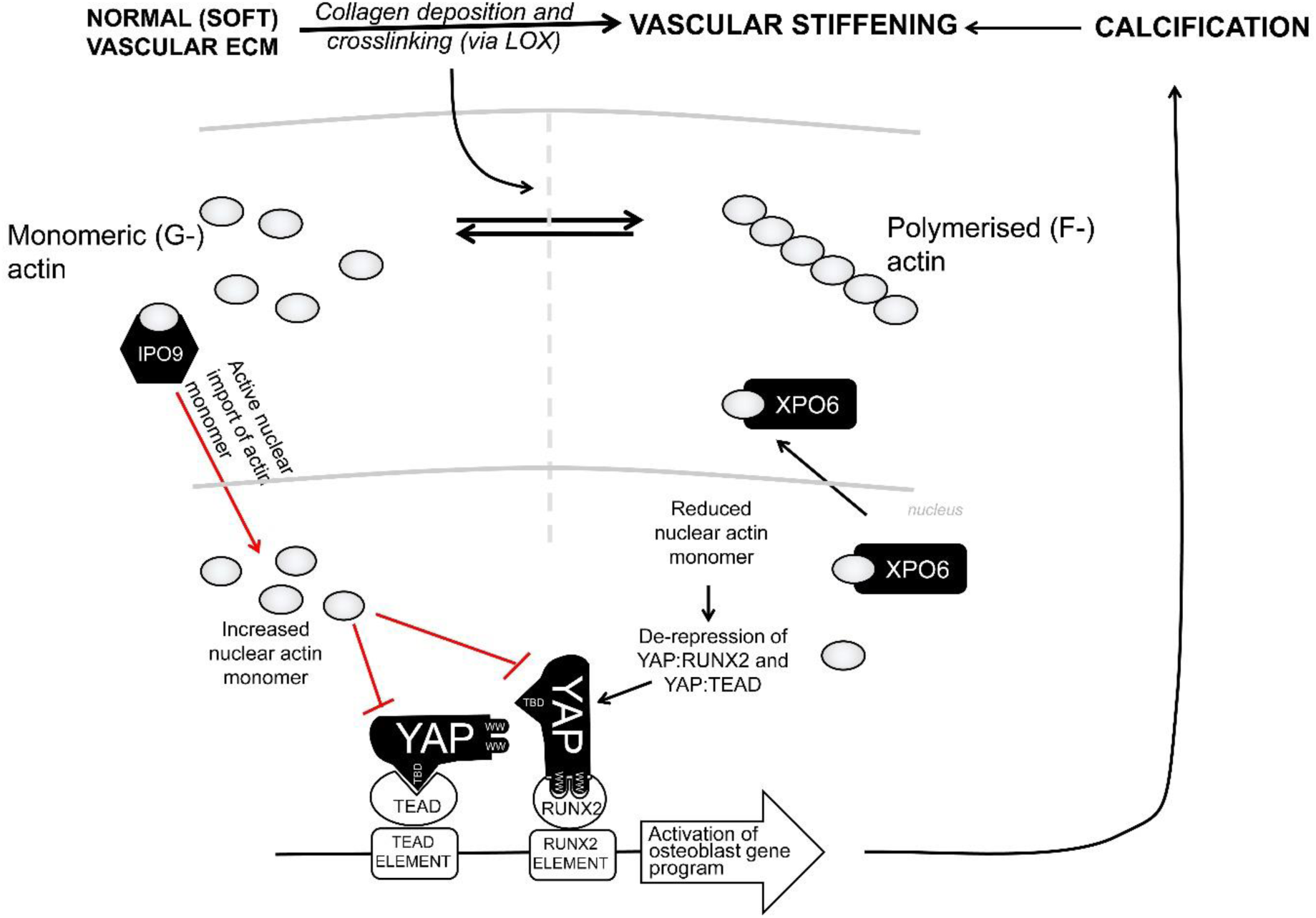

**HIGHLIGHTS:**

- Increased ECM stiffness promotes VSMC calcification.
- Increased ECM stiffness reduces levels of nuclear actin monomer.
- On physiological soft ECM, high levels of nuclear actin monomer inhibits calcification by repressing YAP activation.
- YAP activation promotes calcification by stimulating the activity of TEAD and RUNX2.

## 3.0 INTRODUCTION

Progressive vascular calcification (VC), the pathological deposition of insoluble calcium minerals in the arterial wall, is associated with increased risk of heart disease, stroke and atherosclerotic plaque rupture^1^, with particularly high incidence associated with age, diabetes^2^ and chronic kidney disease^3, 4^, where it is a risk factor for cardiovascular events and all-cause mortality ^5^ ^6^. VC exacerbates progressive vascular stiffening, causing impaired hemodynamics and increased pulse wave velocity (PWV)^7–9^, ultimately resulting in substantial cardiovascular (CV) morbidity and mortality from hypertension, aortic stenosis, cardiac hypertrophy, myocardial and limb ischemia, and heart failure ^10–14^. Despite its prevalence and massive impact, only limited therapeutic options are available. These target the calcification process by management of mineral metabolism but are limited by adverse effects on bone and tooth homeostasis ^15^. There is a clear need for a better understanding of the mechanisms underlying VC, to identify new therapeutic targets to limit its progression.

Despite the significant clinical burden ^16^, our understanding of the molecular mechanisms regulating the development of VC is incomplete. Originally believed to be caused simply by the accumulation of calcium deposits due to dysregulated mineral metabolism, VC is now recognised as a highly regulated process dependent on the transition of VSMC to an osteoblast-like phenotype and activation of a gene expression programme that promotes VC. Expression of osteoblast transcription factors (e.g. *RUNX2, SOX9, MSX2* and *SP7*) and bone related proteins (*BMP2, BMP4, BGLAP, OPG*, *SPARC*, *SPP1* and *ALKP*) is elevated in calcified vessels^17–20^. Furthermore, reduced expression or function of endogenous calcification inhibitors (*ENPP1*, *MGP*, *TNFRSF11B* and *KL*) is also associated with VC^21–23^. VSMC lineage tracing studies in mice with VSMC specific knockout of matrix Gla protein demonstrates that this osteoblastic differentiation occurs *in vivo* prior to deposition of calcium in vessels^24^. This suggests that VSMC adopt an osteogenic phenotype that promotes VC and its associated vascular pathologies. However, the mechanisms regulating acquisition of a VSMC osteogenic phenotype are poorly understood.

Vessel stiffness increases with age as a result of adaptive remodelling, increased collagen deposition, and increased collagen crosslinking^25–27^. Classical CV risk factors, including hypertension, diabetes, dyslipidemia and smoking, all increase vessel stiffness. Importantly, multiple studies have shown that arterial stiffness, determined by PWV, is a predictor for cardiovascular events^28–31^. Moreover, vascular stiffness promotes increased blood pressure, cardiac and vascular fibrosis and inflammation, inducing both vessel and cardiac overload, ultimately leading to atherosclerosis and heart failure^32–34^. Importantly, increased vascular stiffness has been directly linked to modulation of VSMC phenotype, although the role of vascular stiffness in regulating VC is currently unclear^35–37^. However, increased ECM-stiffness is known to enhance osteogenic differentiation of mesenchymal stem cells^38, 39^ suggesting existance a mechanistic link between tissue stiffness and osteogenic phenotype.

Cells sense local tissue stiffness via integrins, which modulate activity of RhoGTPases and actin depolymerising factors such as cofilin, thus relaying information regarding ECM-stiffness to organisation of the actin cytoskeleton^36, 40, 41^. Although the importance of the actin cytoskeleton in VC is currently unclear, several studies have demonstrated that Rho-ROCK inhibition attenuates calcification^42–44^, implicating actin cytoskeleton dysregulation as an important factor in the pathogenesis of VC. The actin cytoskeleton affects cell phenotype and function by modulating the activity of actin-sensitive transcription factors^45^. For example, cytoplasmic:nuclear shuttling of the transcriptional co-factors YAP and TAZ, which bind and activate TEAD transcription factors, is controlled by actin organisation^45^. Importantly, YAP and TAZ have also been implicated in the positive regulation of the osteogenic transcription factor RUNX2. YAP interacts with RUNX2 via its two WW domains and with TEAD via an N-terminal TEAD-binding domain (TBD)^46^. YAP and TAZ have also been implicated in the regulation of calcification. For example, YAP/TAZ promote intramembranous ossification via RUNX2 in endothelial cells^47^ and YAP deletion in osteoblastic lineage cells or mesenchymal stem cells reduces osteogenesis and bone formation^48, 49^.Actin is best known for its function in the cytoplasm as part of the cytoskeleton, where it plays an important role in controlling cell morphology, cell motility and cell signalling. However, actin is also present within the nucleus, where it regulates many essential functions, including nuclear stability, chromatin remodelling and gene transcription. We recently described a novel role of nuclear actin in repressing the expression of genes associated with a synthetic VSMC phenotype^50^, at least in part by antagonising the activity of YAP and TAZ. Here we investigate the role of nuclear actin dynamics in mediating the effects of ECM stiffness on VSMC calcification.

### 4.0 MATERIALS AND METHODS

### 4.1. Reagents

All chemicals were obtained from Sigma unless otherwise stated. Antibodies to YAP (#140745), YAP phospho-serine-127 (#4911), YAP phospho-serine-397 (#13619) were from Cell Signalling Technologies. Antibody to GAPDH (MAB374) was from Millipore. Silencer Select siRNA targeting rat XPO6 (s147550), rat YAP, rat TAZ and non-targeting negative control (#4404020) were from Thermo Fisher Scientific.

### 4.2. Smooth muscle cell culture

Male Sprague Dawley rats were killed by cervical dislocation in accordance with schedule 1 of the U.K. Animals (Scientific Procedures) Act 1986 and Directive 2010/63/EU of the European Parliament and with the approval of the University of Bristol. Human aortic VSMCs (HuVSMCs) were purchased from Promocell and cultured in Smooth Muscle Growth Medium 2 (Promocell). All experiments were replicated for the number of times shown in the text and figures using different batches of cells that were prepared from different animals/donors. Cultures of rat aortic VSMCs (RaVSMCs) were prepared as previously described ^51^ and cultured in Dulbecco’s Modified Eagle’s Medium (DMEM) supplemented with 10% foetal bovine serum, 100 U/ml penicillin/streptomycin and 2.5 mM L-glutamine. Human VSMC were cultured in human smooth muscle cell growth medium 2 (Promocell), supplemented with the supplement pack (Promocell), 100 U/ml penicillin/streptomycin and 2.5 mM L-glutamine. Collagen coated polyacrylamide hydrogels of tuneable stiffness were purchased from Cell Guidance Systems. Unless otherwise states stiff gels had a young’s modulus of 50 kPa and soft gels had a young’s modulus of 1 kPa. Calcification was induced by culture in calcification media (DMEM, 4g/L glucose, 10% FCS, 100 U/ml penicillin/streptomycin and 2.5 mM L-glutamine, 50 µM ascorbic acid, 10 mM β-glycerophophate, 8 mM CaCl_2_).

### 4.3 *Ex vivo* culture of human saphenous vein

Surplus segments of surgically prepared human saphenous vein were obtained from consenting patients undergoing coronary artery bypass grafting (Ethics number REC: 20/NE/0103). Vein segments were dissected in Dulbecco’s Modified Eagle’s Medium (DMEM) supplemented with 100 U/ml penicillin/streptomycin and 2.5 mM L-glutamine. For tensile experiments, vein was opened along its longitudinal axis and transverse segments cut to yield 10 mm segments which were cultured in LOX-conditioned media for 4 days. For calcification assays, vein was cut into ∼1mm rings and cultured in High Glucose DMEM supplemented with 10% foetal bovine serum, 100 U/ml penicillin/streptomycin, and 2.5 mM L-glutamine in the presence of 10 mM β-glycerophosphate, 8 mM CaCl2 and 50 µM ascorbic acid for 7 days. For histology, venous material was fixed in 4% (v/v) formalin/PBS for 20 minutes in preparation for paraffin embedding.

### 4.4 Mechanical analysis of human saphenous vein samples

Human saphenous vein segments were tested in uniaxial tension using a universal testing machine (Instron 3342, Norwood, MA, USA) with a 100 N load cell. Sample thickness and width were measured using a micrometer and a caliper. Uniaxial tensile testing was performed at a constant strain rate of 10 mm·min−1 and up to rupture. Stress-strain curves for all experiments were obtained from axial loading and clamp displacement data recorded during the test. Young’s modulus was calculated by BlueHill software as the ratio between stress and strain in the first linear portion of the stress-strain curve.

### 4.5 Quantichrome calcium assay and lysyl oxidase assay

Extracellular calcium mineral accumulation was quantified using the Quantichrome calcium assay, essentially according to the manufacturers instructions. Briefly, cells or tissue were washed three times in calcium free PBS before extraction overnight in 0.6M HCl. Extracts were centrifuged to remove insoluble material and supernatant assayed using a Promega Discover plate reader. Calcium content was normalised to the wet weight of the tissue. Lysyl oxidase activity in cell conditioned media was quantified using the fluorometric Lysyl oxidase kit (Abcam).

### 4.6 Quantitative RT-PCR and western blotting

Quantification of mRNA and protein levels was performed by RT-qPCR and western blotting respectively, as described previously [<u>24</u>]. Total RNA was extracted using Ambion Pure-Link kits (Thermo Fisher) and was reverse transcribed using QuantiTect RT kit (Qiagen) using random priming. Quantitative PCR was performed using Roche SYBR Green using a Qiagen Roto-Gene Q PCR machine (20’@95 °C;20’@62 °C;20’@72 °C). Primers sequences are described in Supplement Table 1. Data were normalised to total amount of RNA. Western blots were performed using a Mini-Protean II system. Proteins were transferred to PVDF membrane using a semi-dry Turbo blotter system (Bio-Rad) and detected using ECL and a digital ChemiDoc imaging system (Bio-Rad).

### 4.7 Plasmids, siRNA and adenoviral vectors

Replication deficient adenoviral vectors were generated by cloning cDNAs into the multiple cloning site of the pDC515 adenovirus shuttle vector (Microbix) as previously described [<u>36</u>]. Adenovirus expressing FLAG-tagged actin containing a nuclear localisation signal and an R62D mutation to prevent polymerisation was generated by cloning FLAG-NLS-β-ACTIN-_R62D_ cDNA into the EcoR1 and Nhe1 sites of plasmid pDC515 followed by homologous recombination with adenovirus genome vector pBHG-frt in HEK293 cells and has been described previously ^50^. Adenovirus expressing mouse Importin-9 (IPO9) was generated by PCR amplifying the IPO9 cDNA from MGC cDNA clone BC098508 and cloning into the Nhe1 and Sal1 sites of pDC515 ^50^. Adenovirus expressing human Exportin-6 (XPO6) was amplified by PCR and cloned into the Nhe1 and BamH1 sites of plasmid pDC515 ^50^. Adenoviral vectors expressing YAP_S127A_ and TAZ_4SA_ were generated by cloning cDNAs into the multiple cloning site of the pDC515 adenovirus shuttle vector (Microbix) as previously described ^52–54^. YAP mutants of both WW-domains, which disrupts YAP interaction with RUNX2 ^46^, in the context of the YAP_S127A_ activation mutation were generated by mutating W199A, W202A, W258A and P261A by PCR. YAP mutant containing S89A mutation, which disrupts interaction with TEAD ^46^, in the context of YAP_S127A_ was also generated by PCR. Resulting mutants were cloned into pDC515 and recombinant adenoviruses generated by co-transfection of HEK293 cells with the pDC515 expression vector and the adenoviral genome vector pBGHfrt, as previously described ^55^. Clones of replication deficient adenoviruses were amplified by infection of four 150 cm2 flasks of HEK293 cells and the resulting adenovirus purified using AdEasy adenovirus purification kit (Agilent). Purified adenoviral stocks were titrated by end-point dilution plaque assay in HEK293 cells.

Plasmid TEAD-LUC expressing firefly luciferase under the control of a synthetic promoter containing eight copies of a consensus TEAD binding element was obtained from Addgene (#34615) and has been described previously ^54^. RUNX2 reporter plasmid was generated by cloning six copies of the RUNX2 consensus binding element (5’-TGTGGTTT-‘3) upstream of a minimal promoter in vector pNL3.3:secNLUC, encoding a secreted form of nanoluciferase. Additionally, to specifically measure RUNX2 activaion we constructed a GAL4-RUNX2 fusion expression vector, in which the transcriptional activation domain of human RUNX2 was fused to the C-terminus of the GAL4 DNA binding domain. This expression vector was used in conjunction with GAL4-secNLUC nano-luciferase reporter (GAL4-NLUC), which was created by subcloning the 5xGAL4 binding elements from plasmid pG5E1b-LUC (a gift from Ugo Moens, University of Tromsø, Norway) into the Nhe1 and Xho1 sites of pNL3.3[*sec-Nluc*/minP] (Promega). Human lysyl oxidase (LOX) expression vector was produced by PCR amplification of the human LOX cDNA and cloning it into expression vector pDC515, with a C-terminal V5 epitope tag.

### 4.8 Bioinformatics analysis

Gene ontology enrichement analysis was performed using the Gene Ontology Consortium wesite ^56^ ^57^. Geneset enrichment analysis was performed using the GSEA application from the Broad Institute (UC San Diego) ^58^. Motif analysis of differentially regulated gene promoter regions was performed using the FIMO application ^59^. Transcription factor enrichment analysis was performed using the oPOSSUM3 application ^60^.

### 4.9 Transient transfection and reporter gene assays

TEAD reporter gene activity was determined by quantifying the cellular luciferase reporter activity in cells transfected with TEAD-LUC (Addgene #34615). RUNX2 reporter gene activity was determined using plasmid pNL3.3:RUNX2, which expresses secreted nanoluciferase under the control of a promoter containing six RUNX2 binding elements or co-transfection with pNL3.3:GAL4 together with a GAL4-RUNX2 expression vector. Plasmid transfection was performed by electroporation of 1 × 10^6^ cells with up to 5 μg of plasmid DNA using an Amaxa Nucleofector-1.5 (program A-033). For gene silencing, cells were transfected with 100 pmoles of Silencer Select siRNA (Life Technologies). At the indicated times post transfections, cell lysates were prepared in ice cold Cell Culture Lysis (CLB) buffer (Promega) and assayed for firefly luciferase activity using the luciferase assay system (Promega) and a Glomax Discover luminometer (Promega) according to the manufacturer’s instructions.

### 4.10 Quantification of intranuclear actin using DNAse1 staining

Levels of intranuclear actin in fixed cells or *ex vivo* human saphenous vein sections were detected with a DNAse1-Alexa Fluor-594 conjugate, essentially as previously described ^50^. Cells or ex vivo human saphenous vein sections were fixed for 20 min at room temperature in 4% methanol-free formaldehyde and permeabilised with 0.1% Triton-X-100 and blocked in 5% bovine serum albumin for 20 min at room temperature followed by incubation with 9 μg/ml of DNAse1-Alexa Fluor-594 for 20 min at room temperature. Following washing in PBS, nuclei were counter stained with Hoechst 33342. Imaging was conducted using a Leica confocal microscope and the nuclear DNAse1-Alexa Fluor-594 fluorescent signal was quantified in *Z*-slice images passing through the nuclei using an automated ImageJ script.

### 4.11 Quantification of calcification using OsteoSense staining

Paraffin embedded *ex vivo* human saphenous vein sections were deparaffinised and rehydrated by immersion into ethanol solutions (95%, 80%, 70%) for 5 min each. Following a washing step in deionised water staining with Osteosense™680EX (100 pmol/L in PBS) was performed at 37 °C for 24 h, followed by three washing steps in PBS and counter-staining of the nuclei with Hoechst staining, according to the manufacturer’s protocol. Sections were then imaged using an Evident (Olympus) VS200 system with Olympus VS200 ASW software.

### 4.12 Statistical analysis

After testing for Gaussian distribution, statistical analysis was performed using two-way ANOVA, one-way ANOVA with Student-Newman-Keuls post-test or where appropriate a paired student’s t-test, as indicated. *Indicates p < 0.05, **indicates p < 0.01, ***indicates p < 0.001.

### 5.0 RESULTS

### 5.1 VSMC calcification and expression of calcification associated genes is dependent of extracellular matrix stiffness

Increased vascular stiffness is a predictor for cardiovascular events^28–31^. Although vascular calcification is associated with increased vascular stiffness, it is not clear if increased tissue stiffness itself promotes an oestrogenic phenotype and calcification. To test this, we cultured VSMC on collagen coated polyacrylamide hydrogels of tuneable stiffness in the presence of calcification media and quantified VSMC calcification and expression of genes associated with an osteogenic phenotype. VSMC cultured for 7 days on tissue culture plastic in the presence of calcification media resulted in a significant increase in the levels of extracellular acid soluble calcium minerals, compared to cells grown in standard growth media (Figure 1A). Importantly, VSMCs cultured on stiff ECM in calcification media for 7 days deposited significantly more ECM calcification than cells grown on soft ECM, detected by Alizarin red staining (Figure 1B and C) and Quantichrome calcium assay (Figure 1D). Using hydrogels covering a range of physiological stiffnesses from 0.1 to 50 kPa, we demonstrate that VSMC calcification progressively increases as the substrate stiffness increases (Figure 1E).

**Figure 1:**
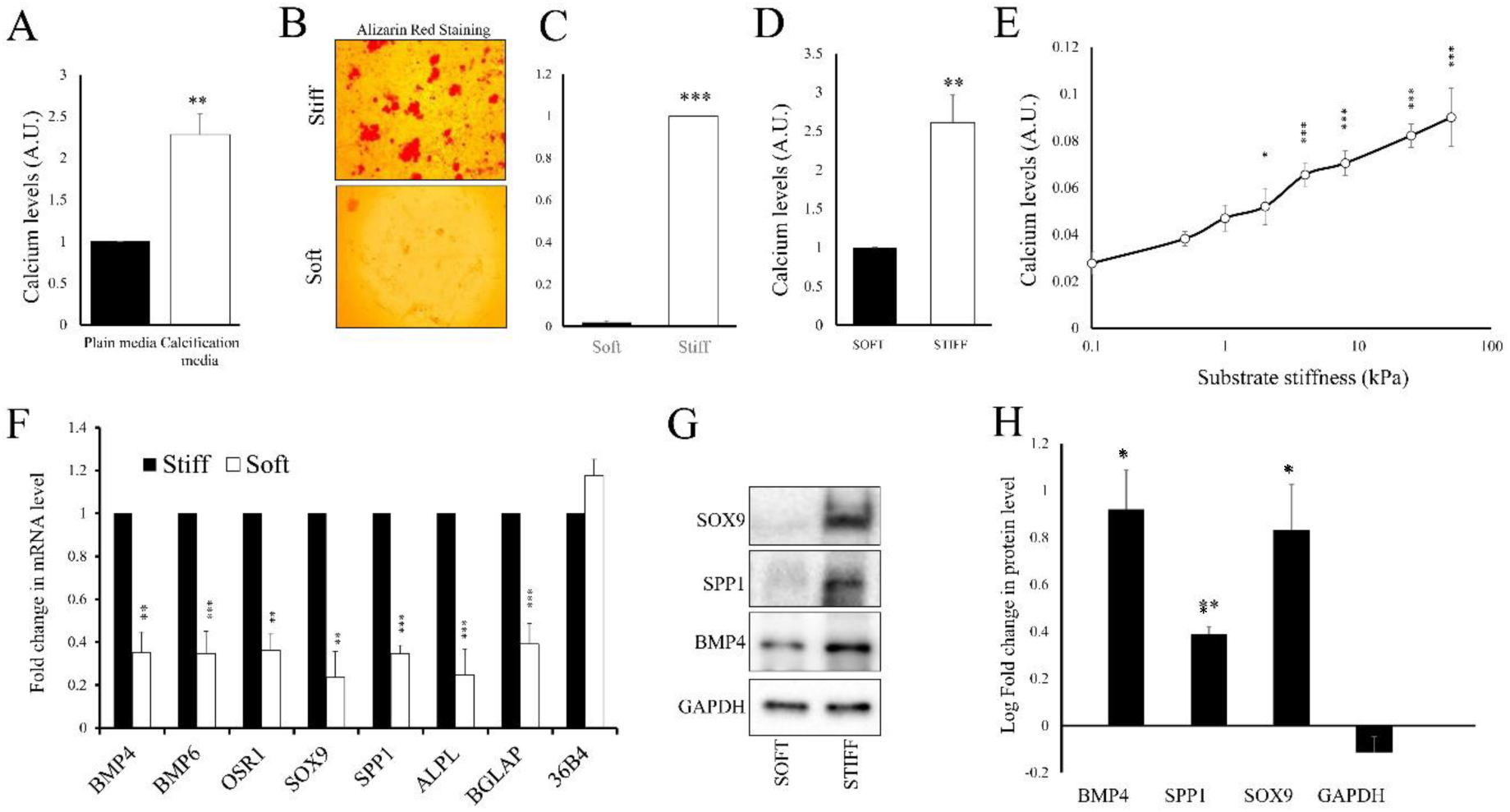
Increased ECM-stiffness promotes VSMC calcification and an osteogenic phenotype Human VSMC were cultured on soft or stiff collagen coated polyacrylamide hydrogels as indicated in normal growth medium or calcification medium for 7 days (A-D). Calcification was quantified using Alazarin red staining (A-C) and quantichrome calcium assay (D). Cells were seeded onto collagen coated polyacrylamide hydrogels of indicated stiffness in calcification media. Calcification was quantified using quantichrome calcium assay (E). Rat VSMC were seeded onto soft or stiff hydrogels. Total RNA was extracted after 24 hours and analysed by RT-qPCR for the indicated mRNAs (F). Total cell lysate was analysed for the indicated proteins by western blotting (G,H). * indicates p<0.05, ** indicated p<0.01, *** indicates p<0.001. Student paired t-test (A, D). One way ANOVA with student Newman Keuls post test (E, F, H).

Analysis of mRNA levels of genes known to be involved in osteogenesis and vascular calcification detected significant increases in the mRNA levels of BMP4, BMP6, OSR1, RUNX2, SOX9, SPP1, ALPL and BGLAP in cells cultured on stiff compared to soft ECM (Figure 1F). Steady state mRNA levels of the housekeeping gene 36B4 were not affected by ECM stiffness. Analysis of protein levels of selected genes (BMP4, SPP1 and SOX9) by Western blotting also detected a significant increase in the levels of these proteins in cells cultured on a stiff ECM compared to cells grown on a soft ECM (Figure1 G and H). Protein levels of the housekeeping gene GAPDH were unaffected.

### 5.2 Increased ECM stiffness reduces levels of nuclear actin monomer

Cells sense the local mechanical properties of the ECM via changes in the organisation of the cytoplasmic actin cytoskeleton, actin polymerisation and actin stress fibre tension. Changes in cytoplasmic actin polymerisation affect the availability of actin monomers that can be actively transported into the nucleus, where they regulate gene expression and VSMC phenotype ^50, 61^. We analysed if ECM stiffness and actin stress fibre tension modulated the levels of nuclear actin monomer in VSMC and if this was an important mechanism in the regulation of VSMC calcification. VSMC cultured on a soft ECM or treated with the myosin-II ATPase inhibitor, Blebbistatin, displayed significantly reduced cell spreading, with a rounded condensed morphology, compared to cells on a stiff ECM or control cells (Figure 2A, D, F and I). Cells on a soft ECM or treated with Blebbistatin, also contained fewer actin stress fibres, compared to cells on stiff ECM or control cells (Figure 2B, G). Importantly, staining with Alexa-fluor 594 labelled DNAse1, which selectively binds to actin monomer ^62^, demonstrated a significant increase in nuclear actin monomer levels in cells grown on a soft ECM or treated with Blebbistatin, compared to cells on a stiff ECM or control cells (Figure 2 C, E, H and J). Taken together, this demonstrates that ECM stiffness and actin stress fibre tension are inversely related to levels of nuclear actin monomer in VSMC.

**Figure 2:**
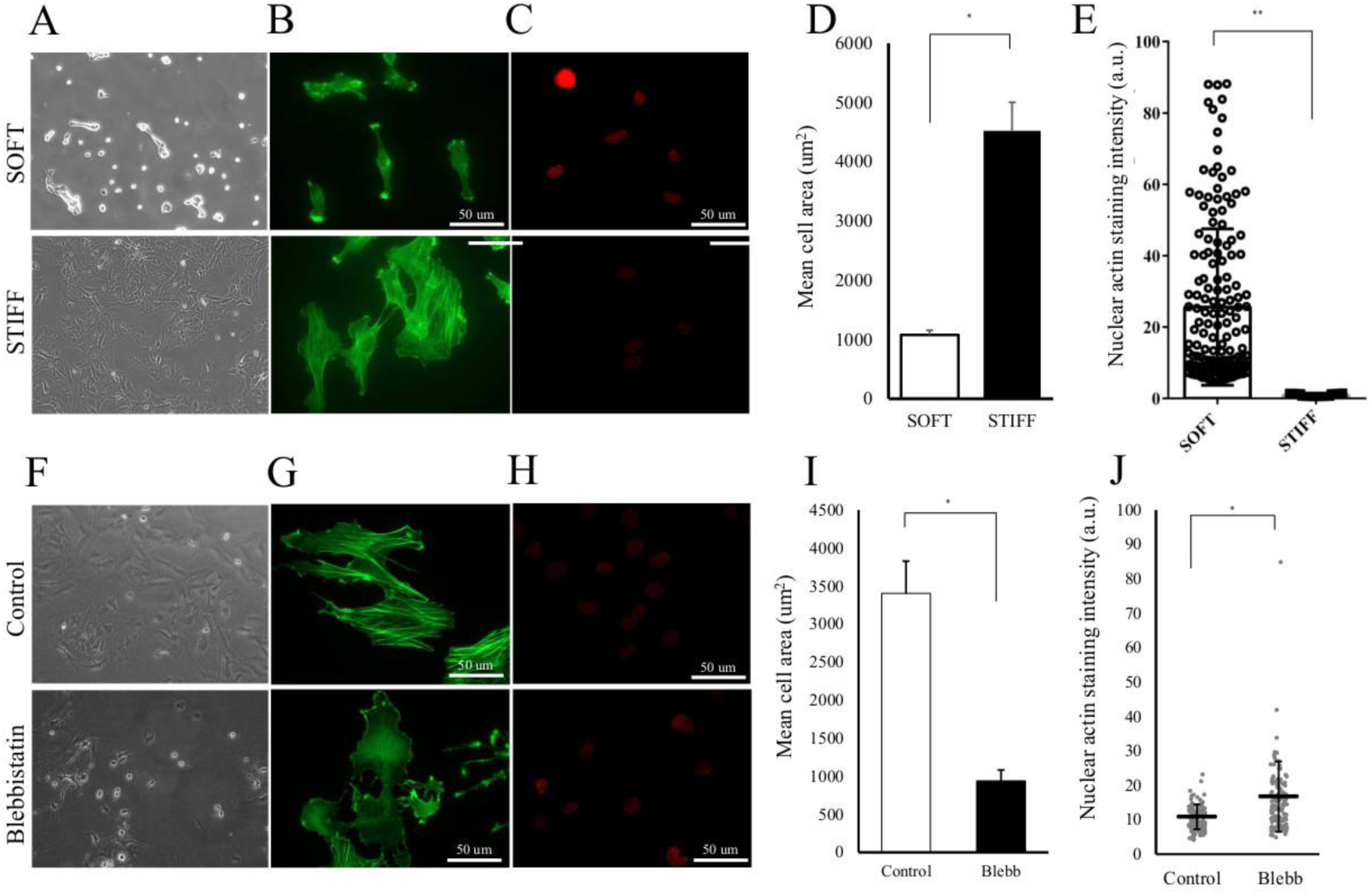
Increased ECM-stiffness promotes cell spreading, actin stress fibre formation and a reduction in nuclear actin monomer levels. VSMC were seeded onto soft or stiff collagen coated polyacrylamide hydrogels (A-E). Cells were analysed by phase contrast microscopy (A), phalloidin staining (B), Alexa-595 DNAse1 staining for nuclear actin mononmer (C, H) and mean cell area (G). Cells were seeded onto tissue culture plastic and analysed by phase contrast microscopy (F), phalloidin staining (G), Alexa-595 DNAse1 staining for nuclear actin mononmer (H, J) and mean cell area (I). * indicates p<0.05, ** indicates p<0.01. Paired student T-test.

### 5.3 Transcriptomic analysis identifies repression of osteogenic genes in VSMC expressing nuclear localised actin monomer

We analysed our previously published transcriptomic dataset ^50^ of genes selectively regulated by expression of a nuclear localised polymerisation-defective actin mutant (NLS-Actin_R62D_) in VSMC. Gene Ontology analysis of NLS-Actin_R62D_ repressed genes (2425 repressed genes in total) identified a significant enrichment of Gene Ontology terms associated with osteoblast differentiation (GO:0001649), Positive regulation of bone mineralisation (GO:0030501), Bone development (GO:0060348), positive regulation of osteoblast differentiation (GO:0045669) and osteoblast proliferation (GO:0033687) (Figure 3A). Gene set enrichment analysis also identified a significant enrichment of an osteogenesis gene set (GO_0001503; FDR<25%) within the nuclear actin repressed genes (Figure 3B). Specifically, RNA-seq analysis detected a significant downregulation of several genes (BMP4, BMP6, OSR1, RUNX2, SOX9 and OSR2) known to be involved in osteogenesis and vascular calcification (Figure 1C). Taken together, this indicates that elevated nuclear actin monomer represses expression of genes associated with an osteogenic phenotype.

**Figure 3:**
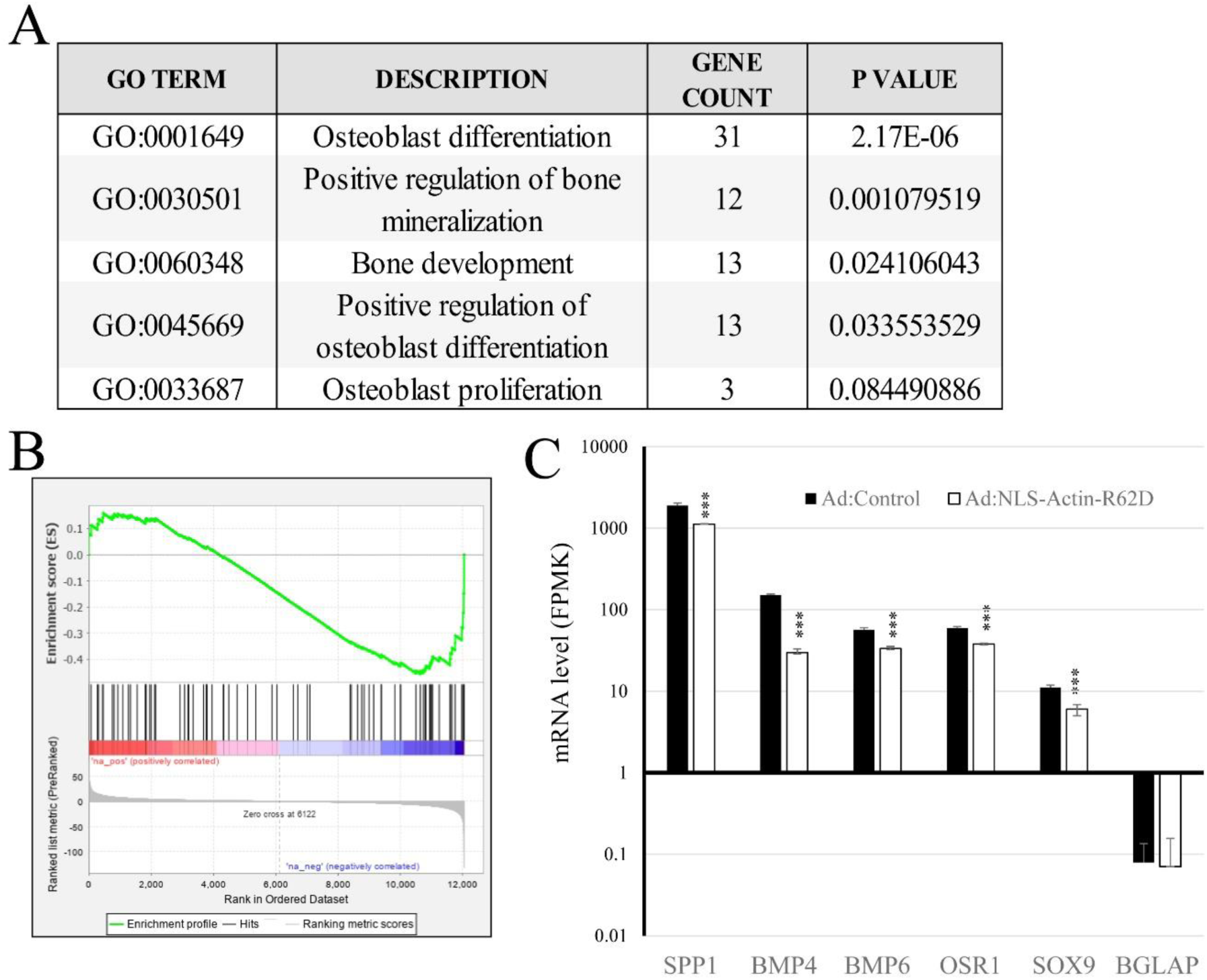
RNA-seq analysis of nuclear actin repressed genes identifies an enrichement of calcification related GO-terms, enrichement of a calcification gene set and repression of calcification associated genes Gene ontology analysis of genes found to by significantly repressed by expression of NLS-Actin_R62D_ (A). Geneset enrichment analysis of NLS-ActinR62D regulated genes with a geneset of genes associated with the bone-mineralisation GO term (GO:0030282). Relative expression of BMP4, BMP6, OSR1, RUNx2, SOX9 and OSR2 mRNA in response to NLS-ActinR62D expression, determined by RNAseq (C)

### 5.4 VSMC calcification and expression of calcification associated genes is repressed by elevated nuclear actin monomer

We initially validated our RNA-seq data using quantitative RT-PCR and extended our analysis to include additional genes known to be involved in tissue calcification (ALPL and BGLAP) ^63^ ^64^. Cultured VSMC were infected with either a control adenoviral vector (Ad:Control) or an adenovirus expressing NLS-Actin_R62D_ (Ad:NLS-Actin_R62D_) and gene expression quantified 24 hours later. Expression of NLS-Actin_R62D_ significantly repressed expression of BMP4, BMP6, OSR1, SOX9, OSR2, SPP1, ALPL and BGLAP (Figure 4A), without affecting mRNA levels of the housekeeping gene 36B4. To further test the hypothesis that elevated nuclear actin represses expression of calcification genes in VSMC, we selectively elevated endogenous nuclear actin monomer levels by overexpressing Importin-9 (IPO9), which actively transports actin monomer into the nucleus ^65^. We previously demonstrated that overexpression of exogenous Importin-9 significantly increases nuclear actin monomer levels in VSMC ^50^. Importantly, Importin-9 (IPO9) expression significantly repressed mRNA expression of BMP4, BMP6, OSR1, SOX9, SPP1 and BGLAP (Figure 4B). Levels of RUNX2, ALPL, BGLAP and the housekeeping gene 36B4 were not significantly affected. Finally, we treated cells with the actin binding drug Latrunculin-B (Lat-B), which inhibits actin polymerisation and increases levels of cytosolic and nuclear actin monomer in VSMC ^50^. Treatment with Lat-B significantly reduced expression of BMP4, BMP6, OSR1, RUNX2, SOX9, ALPL and BGLAP, without affecting 36B4 mRNA levels (Figure 4C). Finally, to test if reduced expression of calcification genes in cells interacting with a soft ECM is, at least in part, mediated by elevated nuclear actin monomer, we over expressed Exportin-6 (XPO6), which has been shown to actively transport actin monomers our of the nucleus and reduce nuclear actin levels in VSMC. In control adenovirus infected cells, mRNA levels of calcification genes were significantly lower on a soft ECM compared to a stiff ECM. Importantly, expression of XPO6, resulted in a significant reduction of calcification gene expression on a soft ECM (Figure 4D)

**Figure 4.**
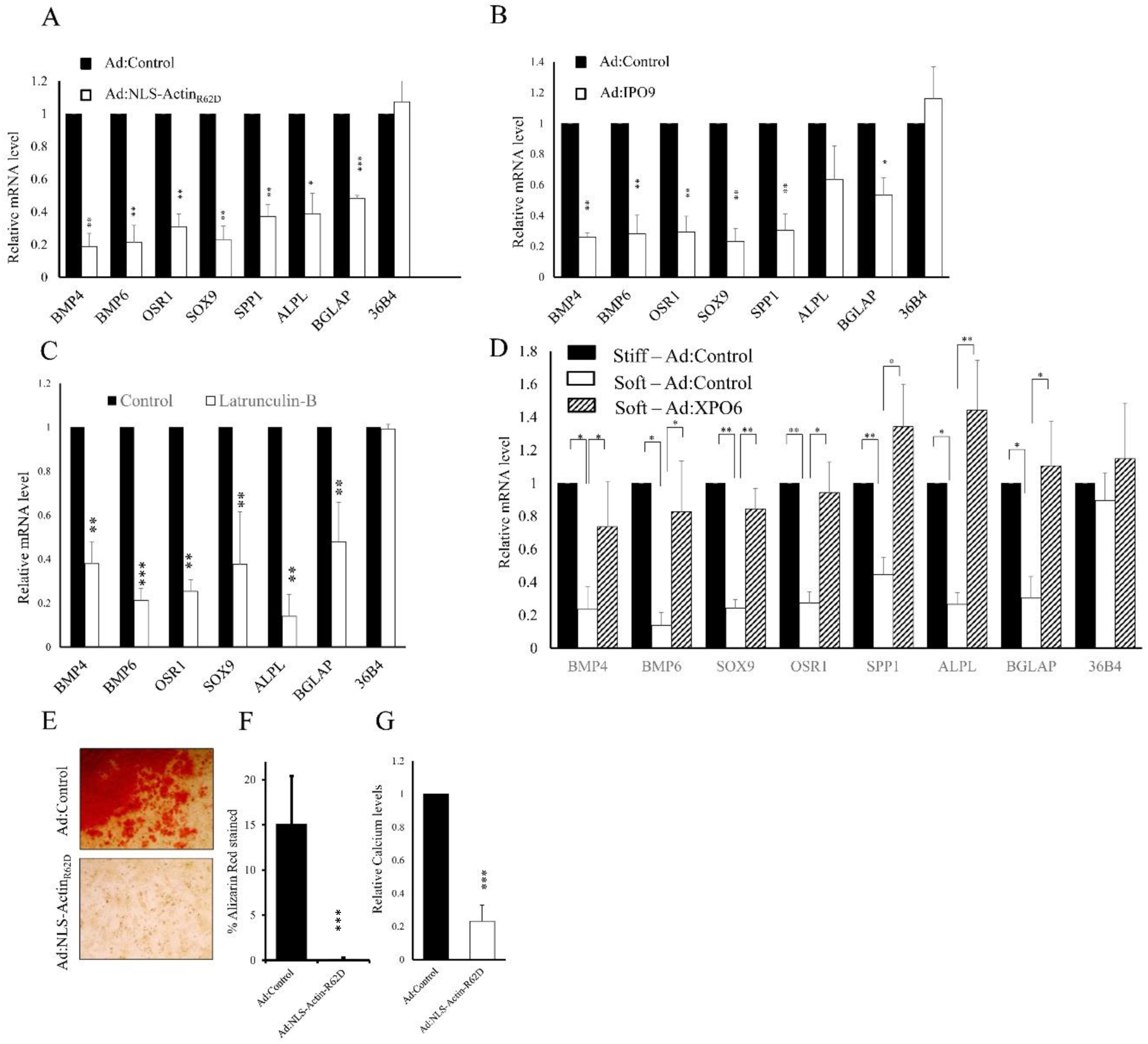
Nuclear actin monomer represses osteogenic gene expression and VSMC calcification Cells were infected with either control adenovirus (Ad:Control), Ad:NLS-ActinR62D (A) or Ad:IPO9 (B), as indicated. Total RNA was analysed for mRNA levels of indicated genes using RT-qPCR. Cells were treated with Latrunculin-B for 6 hours and total RNA analysed for mRNA levels of indicated genes using RT-qPCR (C). Cells infected with Ad:Control or Ad:NLS-ActinR62D were cultured in calcification media for 7 days and calcification quantified using Alizarin red staining (D and E) or quantichrome calcium assay (F). Cells were infected with either Ad:Control or Ad:XPO6 and seeded onto soft or stiff hydrogels. Total RNA was analysed for indicated mRNAs using RT-qPCR (G)

To determine if the observed reduction in the expression of genes associated with calcification, in response elevated nuclear actin monomer results in lower levels of VSMC calcification cells were cultured in calcification media and infected with either Ad:Control or Ad:NLS-Actin_R62D_ adenoviral vectors. Infection with Ad:NLS-Actin_R62D_ completely inhibited VSMC calcification detected by Alizarin red staining (Figure 4E, F) and significantly reduced the levels of ECM calcium, quantified using Quanti-chrome calcium assays (Figure 4G).

### 5.5 Increased tissue stiffness reduces nuclear actin monomer and promotes calcification in human saphenous vein *ex vivo*

To test if modulation of ECM stiffness regulates nuclear actin and calcification in the context of a native vascular ECM, we developed an *ex vivo* model of lysyl oxidase (LOX) induced stiffening of sections of human saphenous vein. LOX is known to cross link collagen fibres and extensive literature documents its role in promoting tissue stiffening ^27, 66–68^. We cloned the human LOX cDNA and over expressed LOX protein, to produce LOX conditioned media. Western blot analysis confirmed the overexpression of the pre-pro LOX in the cell lysate and the processed 28 kDa active LOX fragment in the cell conditioned media (Supplement figure 1A). Elevated LOX activity was confirmed in the LOX conditioned media compared to control media (Supplement figure 1B). Sections of human saphenous vein were incubated in freshly prepared control or LOX conditioned media for 96 hours, with media being replaced after 48 hours. Analysis of vein sections using tensile testing to confirmed that LOX treatment enhanced vein stiffness (Figure 5A). We next used Alexa-594-DNAse1 staining and confocal microscopy, to detect and quantify actin monomer within the nucleus of cells within the control and LOX treated veins. We have previously validated this technique for quantification of nuclear actin levels in cultured VSMC ^50^. To further validate this assay in histological sections of human saphenous vein, we treated vein samples *ex vivo* with the actin binding drugs Lat-B or Jasplaklinolide (to induce actin depolymerisation or polymerisation, respectively). As expected, treatment with Lat-B significantly elevated nuclear actin monomer levels and Jasplakinolide treatment significantly reduced it (Supplement Figure 2A and B). Importantly, LOX-induced vein stiffening was associated with a significant reduction in nuclear levels of actin monomer (Figure 5B). Moreover, LOX-induced stiffening also resulted in a significant increase in vein calcification, which was inhibited by co-incubation with the LOX inhibitor, BAPN (Figure 5C). Furthermore, adenoviral delivery of nuclear targeted polymerisation defective actin (NLS-Actin_R62D_), significantly inhibited LOX-induced vein calcification, detected using either quantichrome calcium assay or staining of histological sections with Osteosense,a bisphosphonate fluorescence dye - specifically stains and quantifies hydroxyapetite crystals (Figure 5D-F). Taken together, these data indicate that stiffening of the native vascular ECM, reduced nuclear actin levels, which is important in permitting the expression of genes required for vessel calcification.

**Figure 5.**
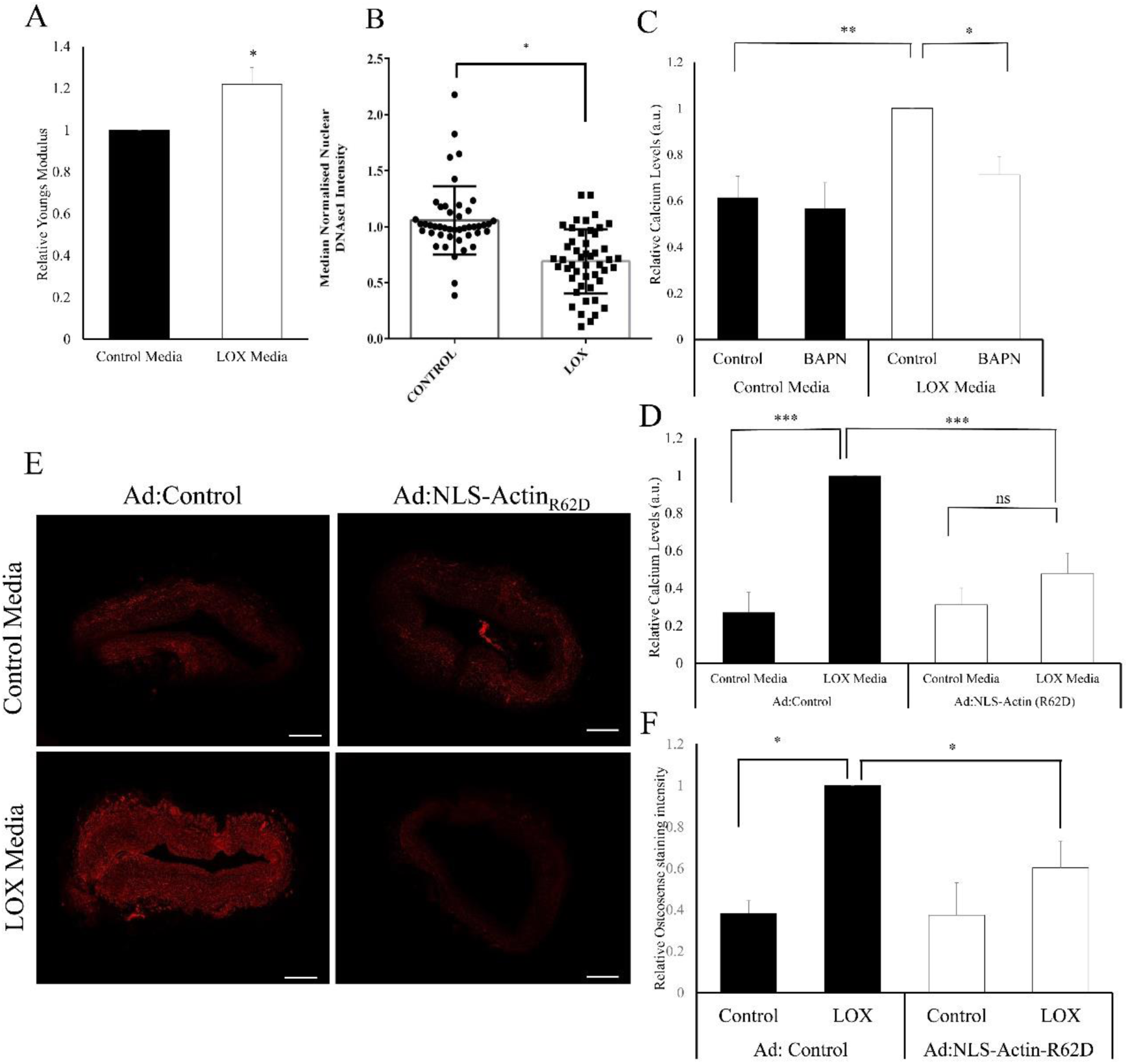
LOX-induced stiffening of human saphenous vein reduces nuclear actin levels and enhances vessel calcification VSMC were transfected with either control plasmid or plasmid expressing human LOX tagged C-terminally with a V5 epitope tag and media conditioned with active LOX collected. Sections of human saphenous vein were incubated with control or LOX conditioned media and effects on tissue stiffness assessed by measuring tissue tensile strength (A). Nuclear actin monomer levels were quantified by staining with Alexa-595 DNAse 1 and confocal microscopy (B). Vein was treated with control of LOX conditioned media in the presence or absence of the LOX inhibitor BAPN and calcification quantified 7 days later using quantifchome calcium assay (C). Vein was treated with control or LOX conditioned media followed by infected with Ad:Control or Ad:NLS-ActinR62D adenoviruses. Calcification was quantified after 7 daysusing quantifchome calcium assay (D) or histological sections stained with ostrosense (E and F)

### 5.6 ECM stiffness and nuclear actin dependent regulation of RUNX2 and TEAD controls VSMC calcification

We analysed the promoter sequences of all nuclear actin repressed genes to identify enrichment of binding elements of transcription factors that may be involved in regulating VSMC calcification in response to changes in nuclear actin levels and ECM stiffness. Using oPOSSUM3^60^ we identified enrichment of TEAD and RUNX2 binding elements in the promoters of genes repressed >1.5 fold by nuclear actin monomer (Figure 6A). Interestingly, genes associated with VSMC calcification (RUNX2, BMP2, BMP3, BMP4, BMP6, OSR1, OSR2, SOX9, ALPL, SPP1 and BGLAP contained multiple TEAD and RUNX2 elements between the TSS and -10kbp, that were often in close proximity (Figure 6B). This implies that nuclear actin monomer-dependent repression of these genes may be mediated, at least in part, via inhibition of TEAD and RUNX2 activity. We therefore quantified the effect of increased nuclear actin monomer levels on TEAD and RUNX2 reporter gene activity. Exogenous expression of NLS-Actin_R62D_ significantly reduced reporter gene activity under the control of synthetic promoters containing multimerised copies of either TEAD or RUNX2 binding elements (Figure 6C). Similar inhibition was observed when cells were treated with the actin depolymerising agent, Lat-B. Since we observed an inverse relationship between ECM stiffness and levels of actin monomers within the nucleus, we next quantified TEAD and RUNX2 activity in cells grown on stiff or soft substrates. This demonstrated significantly greater TEAD and RUNX2 reporter gene activity in cells grown on stiff ECM compared to soft substrates (Figure 6D and E). To further test that reduced activity of TEAD and RUNX2 in cells interacting with a soft ECM was, at least in part due to the elevated of actin monomer within the nucleus of these cells, we overexpressed XPO6 to prevent nuclear actin monomer accumulation and rescue TEAD and RUNX2 activity. In cells transient transfected with a control plasmid, TEAD (Figure 6F) and RUNX2 (Figure 6G) activity was significantly lower in cells interacting with a soft compared to a stiff ECM. Exogenous expression of XPO6 resulted in a small and non-significant trend towards increased TEAD and RUNX2 activity. However, exogenous expression of XPO6 resulted in a significant increase in reporter gene activity in cells interacting with a soft ECM (Figure 6 F,G). This is consistent with at least a partial role of nuclear actin monomer in the repression of TEAD and RUNX2 activity in cells interacting with soft matrices.

**Figure 6:**
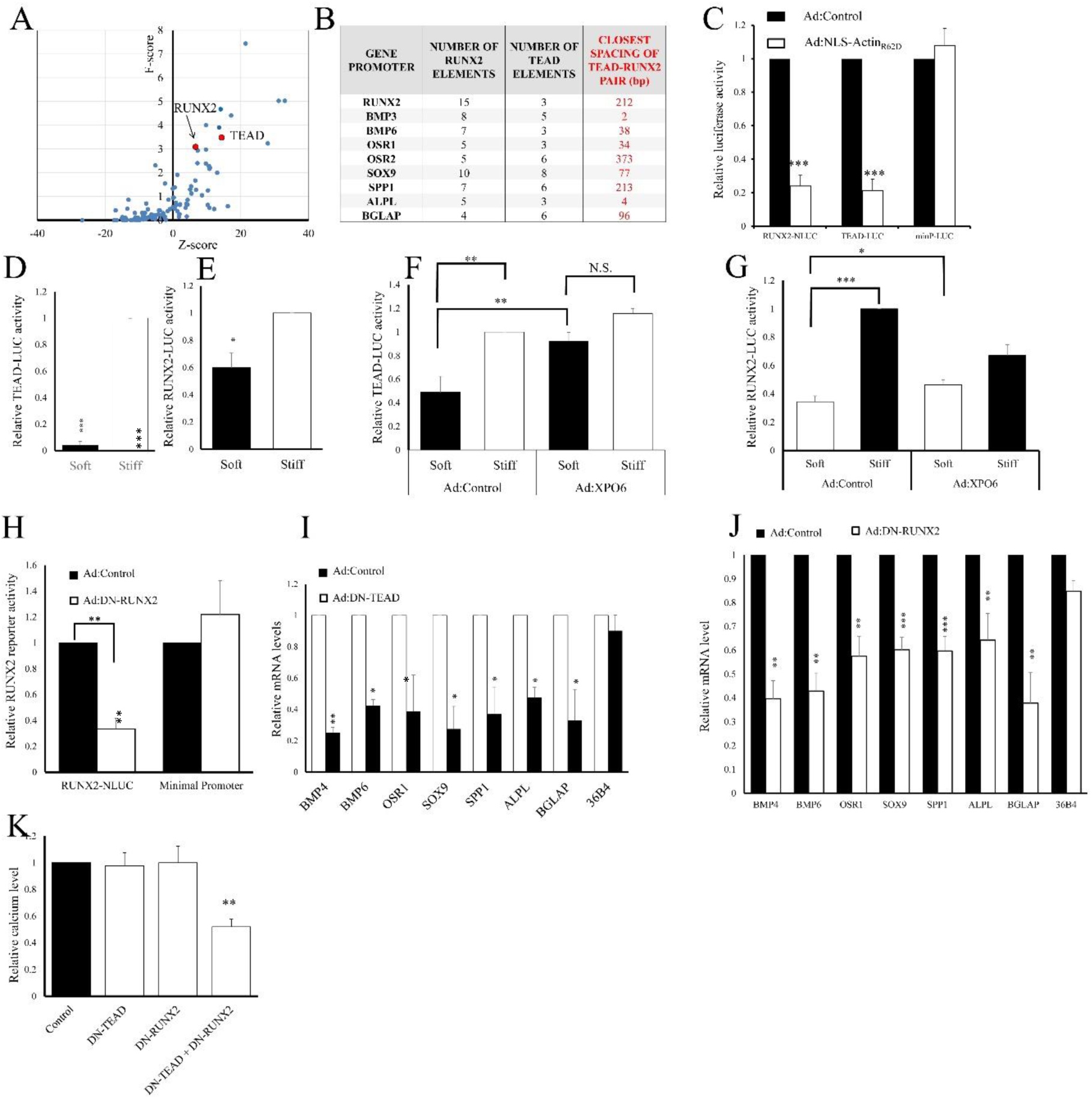
Nuclear actin monomer mediates the effects of ECM stiffness and inhibits TEAD and RUNX2 activity Transcription factor enrichment analysis of the promoter regions (-10kbp to TSS) of NLS-ActinR62D repressed genes (A) Table summarising total number of consensus RUNX2 or TEAD binding elements in promoter regions of indicated genes and minimum distance between elements. VSMC were transfected with TEAD, RUNX2 or control luciferase reporter plasmids and infected with Ad:Control or Ad:NLS-ActinR62D adenviruses (C). Cells were transfected with TEAD (D) or RUNX2 (E) reporter plasmids and seeded onto soft or stiff hydrogels. Cells were transfected with TEAD (F) or RUNX2 (G) reporter plasmids and infected with Ad:Control or Ad:XPO6 before seeding onto soft or stiff hydrogels. Cells were transfected with RUNX2 reporter plasmid and infected with Ad:Control or Ad:DN-RUNX2 (G). Reporter activity was quantified after 24 hours (C-G). Cells were infected with Ad:DN-RUNX2 (H) or Ad:DN-TEAD (I) and total RNA analysed for levels of indicated mRNAs using RT-qPCR.

We next tested the functional importance of TEAD and RUNX2 activity for VSMC calcification. We employed dominant-negative mutants of TEAD and RUNX2, constructed by fusing the relevant DNA binding domain to the transcriptional repressor domain of Engrailled. We previously demonstrated that expression of DN-TEAD, strongly represses TEAD activity ^53, 69^. Expression of DN-RUNX2, also strongly repressed RUNX2 reporter activity (Figure 6H). Expression of either DN-TEAD (Figure 6I) or DN-RUNX2 (Figure 6J) significantly repressed the mRNA levels of the calcification genes (BMP4, BMP6, OSR1, SOX9, SPP1, ALPL and BGLAP). Quantification of ECM calcium using Quantichrome assays demonstrated that, DN-TEAD or DN-RUNX2 alone had no effect on VSMC calcification. However, dual inhibition of TEAD and RUNX2 significantly reduced calcium levels (Figure 6K).

### 5.7 YAP mediates the stiffness-dependent activation of TEAD and RUNX2 and VSMC calcification

YAP has previously been described as a mechano-sensitive co-factor for TEAD family transcription factors ^45^. Importantly, YAP has also been reported to regulate the activity of RUNX2, although the precise nature of this regulation is unclear, with some groups reporting YAP-mediated activation of RUNX2 and others reporting repression ^46, 70–72^. We therefore investigated the role of YAP, in the stiffness dependent regulation of both TEAD and RUNX2 and the role of these two pathways in VSMC calcification. We initially quantified YAP phosphorylation, a surrogate marker of YAP inactivation. We detected a significant reduction in the levels of YAP phosphorylation at Y397, a marker of YAP inactivation, in cells cultured on a stiff ECM (Figure 7A). This is consistent with elevated YAP activation in cells on a stiff ECM.

We next overexpressed constitutively active mutants of YAP, and the related co-factor TAZ, and quantified effects on TEAD and RUNX2 activity. Forced expression of either YAP or TAZ significantly activated TEAD reporter activity (Figure 7 B), consistent with the previously reported TEAD co-factor activity of these proteins ^73^. Importantly, only expression of YAP, but not TAZ, activated RUNX2 reporter gene activity (Figure 7C). To further confirm this selectivity of RUNX2 for YAP over TAZ, we performed siRNA-mediated gene silencing. As expected, silencing of YAP or TAZ individually, significantly reduced TEAD activity, and dual silencing resulted in a greater inhibition than achieved when silenced separately (Figure 7D). Importantly, only silencing of YAP, but not TAZ, repressed RUNX2 activity (Figure 7 E). Dual silencing had a similar effect of YAP silencing alone. Together, these data demonstrate that while YAP and TAZ can both regulate TEAD, only YAP controls RUNX2 activity in VSMC.

**Figure 7:**
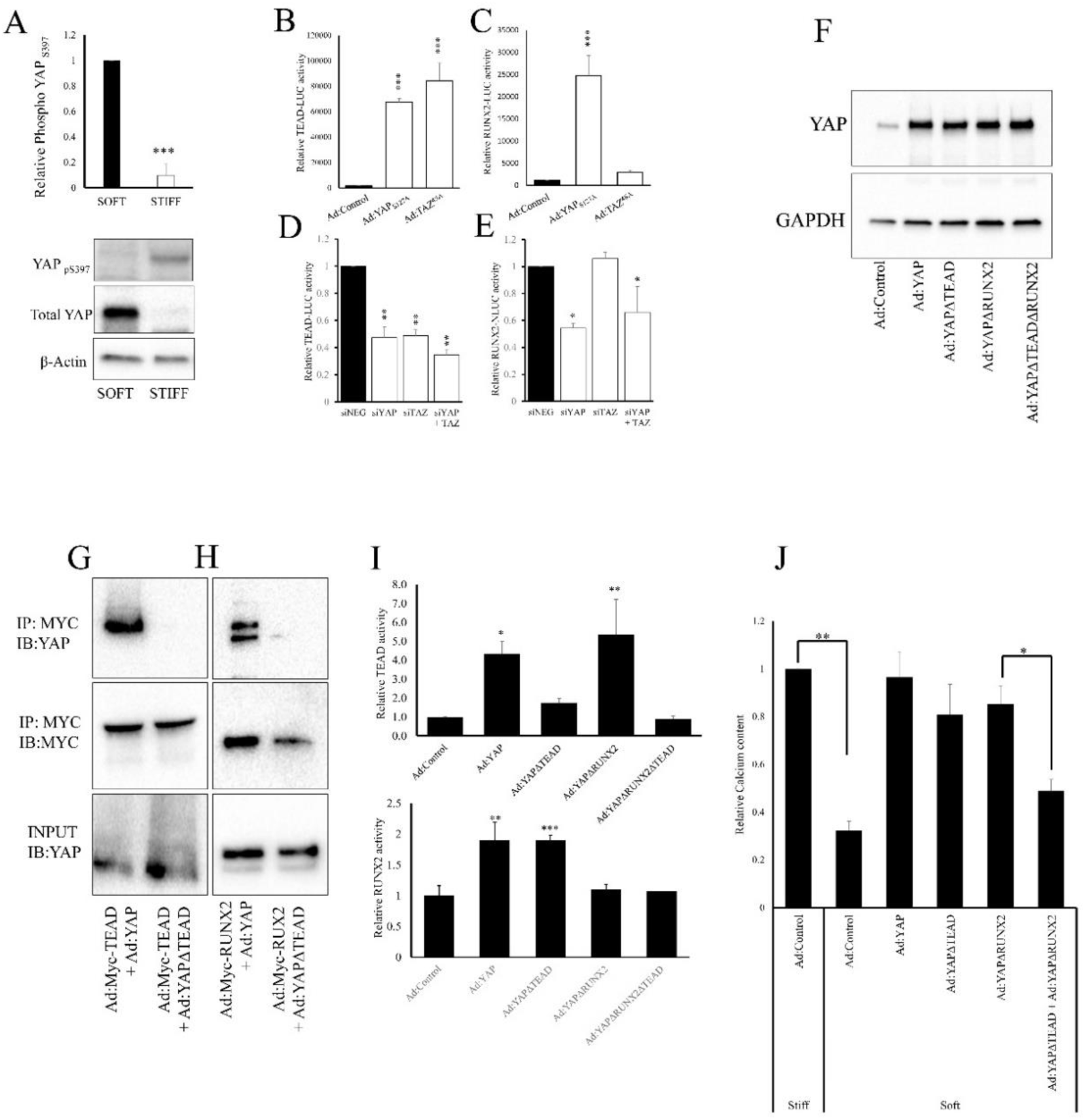
YAP mediates ECM-stiffness-dependent calcification via both TEAD and RUNX2. VSMC were seeded onto soft or stiff hydrogels and total cell lysate analysed for YAP pS397, total YAP and β-actin by western blotting (A). Cells were transfected with TEAD (B) or RUNX2 (C) reporter plasmids followed by infection with Ad:Control, Ad:YAP or Ad:TAZ. Cells were transfected with siRNA targeting YAP, TAZ or YAP plus TAZ, together with reporter plasmids for TEAD (D) or RUNX2 (E). Luciferase activity was quantified after 24 hours. Cells were infected with either Ad:Control, Ad:YAP, Ad:YAPΔTEAD, Ad:YAPΔRUNX2 or Ad:YAPΔTEADΔRUNX2. YAP protein levels were assessed after 24 hours by western blotting (F). Cells were transfected with plasmid expressing either Myc-TEAD or Myc-RUNX2. Cells were also infected with Ad:YAP, Ad:YAPΔTEAD or Ad:YAPΔRUNX2, as indicated. YAP-TEAD (G) or YAP-RUNX2 were detected by Myc-IP and western blotting for coimmunoprecipitated YAP. Cells were transfected with TEAD (I) or RUNX2 (J) reporter plasmids and infected with the indicated YAP mutant adenoviral vector. Cells were seeded onto soft or stiff hydrogels and infected with either Ad:Control or the indicated YAP mutant adenovirus. Effects of calcification were quantified using quantichrome calcium assay (K)

To further test the importance of YAP-dependent activation of both TEAD and RUNX2 for stiffness induced VSMC calcification, we generated a series of YAP mutants based on the well characterised Y397A activation mutation. We first generated a mutated YAP (S94A) defective in TEAD binding (referred to as YAPΔTEAD herein for simplicity). We also generated a version of YAP containing two mutations in each of the two WW domain (T199A/P202A and T258/P261A), previously shown to be essential for RUNX2 interaction (referred to a YAPΔRUNX2. Lastly, we generated a double mutant, defective in binding to both TEAD and RUNX2 (S94A+ T199A/P202A and T258/P261A; referred to as YAPΔTEADΔRUNX2). Equal expression of these exogenous YAP mutants was validated as even by Western blotting (Figure 7F). Co-immunoprecipitation analysis confirmed that the YAPΔTEAD (Figure 7G) and YAPΔRUNX2 (Figure 7H) mutations disrupted interaction with TEAD and RUNX2 respectively. Reporter gene analysis demonstrated with active-YAP (S127A) and YAPΔRUNX2 significantly enhanced TEAD activity (Figure 7I). However, the YAPΔTEAD mutation, either alone or in combination with the RUNX2 binding domain mutations i.e. YAPΔTEADΔ RUNX2 were unable to enhance TEAD activity (Figure 7I). Conversely, active YAP and YAPΔTEAD were able to stimulate RUNX2 activity (Figure 7J), but YAPΔRUNX2 or YAPΔRUNX2ΔTEAD were not (Figure 7J). Finally, we tested the ability of these mutants to promote VSMC calcification in cells interacting with a soft ECM. Consistent with our earlier experiments, cells on a stiff ECM displayed significantly greater calcification compared to those on a soft ECM (Figure 7K). Exogenous expression of YAP (competent in both TEAD and RUNX2 activation) enhanced calcification of cells on a soft ECM to levels equal to that in cells interacting with a stiff ECM (Figure 7K). Likewise, expression of YAPΔTEAD or YAPΔRUNX2 also rescued calcification to levels that were not statistically different from control cells on a stiff ECM. Importantly, expression of YAPΔTEADΔRUNX2 only rescued calcification to levels that were significantly lower than that observed in control cells on a stiff ECM (Figure 7K). Taken together, these data demonstrate that ECM stiffness promotes YAP-dependent activation of TEAD and RUNX2 and that both pathways are required for maximal VSMC calcification.

### 6.0 DISCUSSION

In this study we present evience supporting the existence of a novel mechanism that regulates the adoption of a VSMC osteogenic phenotype in response to changes in the stiffness of the extracellular matrix. We show that increased ECM-stiffness reduces levels of actin monomer within the nucleus. This reduction in nuclear actin derepresses YAP, which promotes an osteogenic phenotype by activating both YAP-TEAD and YAP-RUNX2 transcriptional complexes (see schematic representation Figure 8).

**Figure 8:**
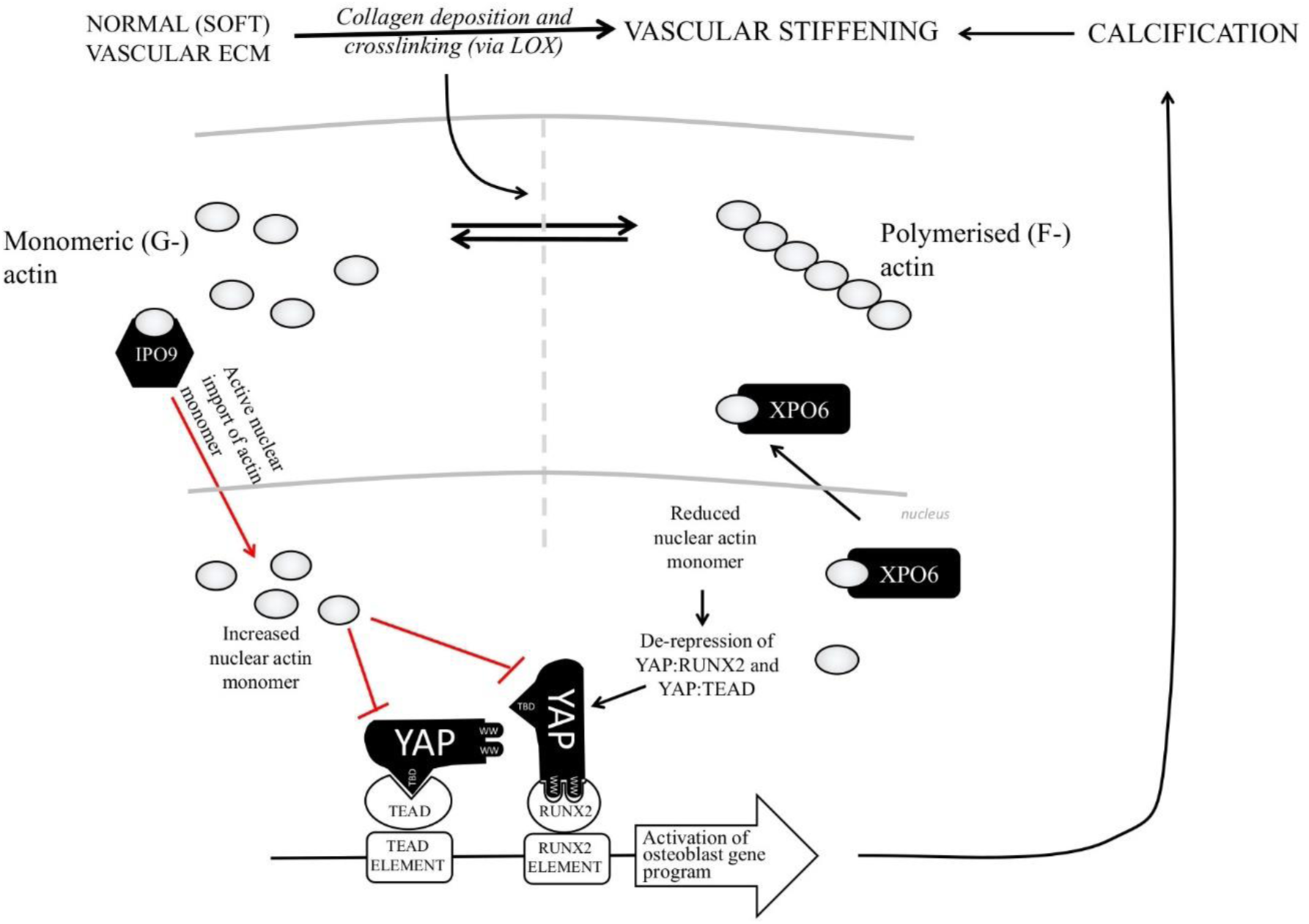
Schematic representation of the mechanisms controlling VSMC calcification in response to ECM stiffness. Increased ECM stiffness induced cytoplasmic actin polymerisation and reduces the levels of actin monomer in the nucleus. Nuclear actin monomer represses YAP activity. Hence the reduction of nuclear actin monomer in response to increased ECM-stiffness results in de-repression of YAP. YAP promotes a VSMC osteogenic phenotype by activating TEAD and RUNX2.

Vascular stiffening is a hallmark of the ageing process and has been associated with a number of diseases, including atherosclerosis, diabetes and chronic kidney disease ^74, 75^. Age-related vascular stiffening is believe to occur as a result of decreased elastin production and an increase in elastin breakdown, as well as increased collagen deposition and subsequent collagen cross linking, via the action of enzymes such as LOX and PLOD ^76–78^. Arterial stiffening is an early marker for increased cardiovascular risk, myocardial infarction and heart failure ^79^. The immediate and direct effects of increased vascular stiffness include hypertension, elevated end systolic pressure, cardiac overload and ultimately cardiac fibrosis and heart failure. At the cellular level, resident vascular smooth muscle cells sense and respond to changes in the compliance of the local ECM by altering their phenotype. For example, VSMC cultured on stiff ECM have lower expression of contractile genes, considered markers of the classical differentiated phenotype, compared to VSMC cultured on softer ECM ^35^. At the same time, expression of genes associated with increased proliferation is elevated in VSMC on stiff ECM ^80^. This suggests that mechanically induced phenotypic changes in VSMC contribute towards vascular remodelling and creation of a vicious cycle of vascular stiffening ^81^

In this study, we highlight for the first time the role of nuclear actin dynamics in mediating mechano-sensitive VSMC calcification. The existence of actin within the nucleus was first reported many years ago ^82–84^. Despite this early research, it is only recently that we are beginning to appreciate the functional importance of nuclear actin and how it is modulated by pathophysiolgocal signals. Actin within the nucleus is now known to play a central role in the regulation of RNA polymerase activity, gene transcription, chromatin remodelling and regulation of cell fate decisions, including apoptotic cell death and differentiation ^85^. In VSMC, we recently demonstrated that elevated nuclear actin repressed cellular processes associated with phenotypically modulated VSMC, including proliferation, migration ^50^ and adoption of a pro-inflammatory phenotype ^86^. Furthermore, we demonstrated that nuclear actin levels were responsive to and mediate the effects of physiological cAMP signalling ^50, 86^. Here we extend these findings, showing that nuclear actin levels in VSMC are responsive to changes in the local ECM stiffness. Furthermore, we show that these changes are important in the regulation of genes associated with an osteogenic phenotype and ultimately, VSMC calcification. The mechansism underlying how changes in ECM compliance result alters nuclear actin momoner levels is currently unknown. However, our data here and previously published ^50, 86^ indicate that nuclear actin monomer levels reflect, at least to some extent, the polymerisation status of the cytoplasmic actin cytoskeleton. For example, we recently demonstrated that cAMP signalling induces a reduction of cytosolic polymerised F-actin, an increase in cytosolic actin monomer (G-actin) and a concomitant increase in nuclear actin. Here we show that increased ECM stiffness promotes cytosolic F-actin polymerisation and is associated with a reduction in nuclear actin monomer levels. This suggests that the cytosolic actin cytoskeleton acts as an environmental sensor, including acting as a biomechanical sensor, and the downstream changes in nuclear actin monomer levels are critical in mediating transcriptional changes that ultimately modulate cellular phenotype. However, it is unlikely that nuclear actin monomer levels simply mirror cytosolic levels, due to passive diffusion into the nucleus. A complex of proteins, including importin-9 (IPO9), cofilin and RAN_GDP_, have been implicated in actively transporting actin monomers into the nucleus^65^, whereas a complex of Exportin 6 (XPO6). profilin and RAN_GTP_ actively transport actin out of the nucleus ^87^. Our data shows that overexpression or silencing of Exportin 6 or Importin-9 effectively uncouples the nuclear actin level from the cytoplasmic actin pool.

A large body of literature documents how nuclear actin controls gene expression ^50, 85, 86, 88–90^. For example, several studies have shown that actin can interact with several chromatin remodelling and histone modifying complexes ^91^. Nuclear actin has also been shown to directly interact with the SRF cofactors, MRTF-1 and MRTF-2, via their RPEL domains, promoting MFTF1/2 nuclear export and repression of SRF-dependent transcription ^92^. We recently demonstrated that nuclear actin mediated the suppression of YAP-TEAD activity and that this was responsible for the anti-mitogenic effects of cAMP in VSMC ^50^. The role of YAP in mechanotransduction is well documented ^45, 93^. Core Hippo pathway kinases are activated by mechanical cues, which cause a loss of tension in, or depolymerisation of, the cytoplasmic actin cyctoskeleton, triggering the phosphorylation and inactivation of YAP. However, our recent data also implicates nuclear actin in the suppression of YAP function ^50^. For example, modulation of nuclear actin levels by either expression of NLS-Actin_R62D_ or overexpression of XPO6 or IPO9 also modulated YAP-TEAD activity in a manner inversely related to nuclear actin levels. Here we show that YAP-TEAD activity is enhanced in cells interacting with a stiff ECM, where nuclear actin monomer levels are reduced. We also present data demonstrating that ECM-stiffness-dependent activation of YAP promotes an osteogenic gene expression program and VSMC calcification. Interestingly, genetic deletion of YAP, together with its paralog TAZ, results in upregulation of osteogenic genes and increased calcification^94^. This likely reflects the loss of cytosolic YAP, which has been implicated in sequestering signalling proteins and preventing their translocation into the nucleus ^94^. Our data supports a role of active nuclear YAP in the promotion of VSMC calcification. For example, using active mutants of YAP, which display constitutive nuclear localisation, we showed that active YAP can drive VSMC calcification, even in cells interacting with a soft ECM. Furthermore, inhibition of TEAD, the best characerised nuclear target of YAP, inhibits osteogenic gene expression.

It should be noted that our TEAD reporter contains multiple copies of a TEAD consensus binding element and measures pan-TEAD (TEAD 1-4) activity. Therefore, we are unable to determine, which of the TEAD family members are being modulated in our experiments. Our dominant negative TEAD is based on the TEAD1 DNA binding domain and so may have a selectivity for TEAD1 inhibition. This is important as TEAD4 has been shown to inhibit VSMC calcification ^95^. TEAD family members are known to have divergent functions ^96^, so it is likely that TEAD 1-3 are involved in mediating ECM-stiffness-dependent calcification. Determining the role of specific TEAD family members in calcification should be a focus of future research. Nethertheless, our data establishes that ECM stifffness-induced YAP activation enhances TEAD-dependent transcription, which contributes towards acquisition of an osteogenic phenotype (see Figure 8). Our new data also highlights the involvement of second YAP-dependent pathway centered around activation of RUNX2. Although TEAD family transcription factors are the most extensively characterised YAP binding partners ^97–100^, RUNX2 was the first YAP interaction partner identified ^70^. However, the functional importance of the YAP-RUNX2 interaction is poorly understood and less well characterised than the YAP-TEAD interaction. Interestingly, YAP has been reported to act as both an activator ^46^ and a repressor ^72^ of RUNX2 activity. Our data, demonstrates that YAP can act as a mechano-sensitive activator of RUNX2. It is notable that our data demonstrates that TAZ did not display any ability to activate RUNX2, despite being equally effective as YAP in mediating TEAD activation and containing WW-domains implicated in RUNX2 interaction ^72^. Activation of RUNX2 by YAP appears to be direct since we were able to co-imunoprecipitate YAP with RUNX2; an interaction that was ablated by mutation of the YAP WW-domains, previously implicated in mediating interaction with RUNX2 ^72^. Since YAP enhances RUNX2 activity in VSMC, we propose that the inhibitory effects of nuclear actin on RUNX2 are mediated via YAP, involving a mechanism that is common to both RUNX2 and TEAD.

The role of RUNX2 in promoting osteogenic gene expression is well documented ^101–103^. However, our data indicates that RUNX2 activation alone is not sufficient for VSMC calcification. For example, inhibition of RUNX2 activitity, using a dominant negative RUNX2 fusion protein, failed to inhibit calcification. Similarly, TEAD inhibition alone, was inefffective. Our data indicates that coordinated TEAD and RUNX2 activation is required for VSMC calcification. Cooperation between TEAD and RUNX2 transcription factors has been reported previously. For example, activation of both TEAD and RUNX2 is essential for maximal proliferation of cardiac fibroblasts ^104^ and ChIP-seq analysis identifies an enrichment of TEAD-binding elements within RUNX2 ChIP-seq peaks ^105^, implying a conserved functional interaction. This may mean that pharmacological inhibition of either TEAD or RUNX2 may not be effective in limiting calcification. Our data suggests that targeting YAP activation may be more effective, as this would reduce both TEAD and RUNX2 activity. An alternative approach may be to pharmacologically manipulate nuclear actin monomer levels, as this acts as a key biomechanical sensor that interprets ECM-stiffness signals and controls osteogenic differentiation of VSMC.

In summary, we show that ECM stiffness controls VSMC calcification by reducing levels of actin monomer within the nucleus. This derepresses YAP, which activates both TEAD and RUNX2 to promote osteogenic gene expression. This suggests that YAP and nuclear actin may be valuable targets for the pharmacological inhibition of vascular calcification.

## 7.1 ACKNOWLEDGMENTS

We acknowledge the technical support and assistance provided by Dr Katy Jepson at the Wolfson Bioimaging facility, University of Bristol

## 7.2 SOURCES OF FUNDING

This work was funded by British Heart Foundation research project number PG/20/10292

## 7.3 DISCLOSURES

None

## SUPPLEMENTARY MATERIAL

**Figure 1:**
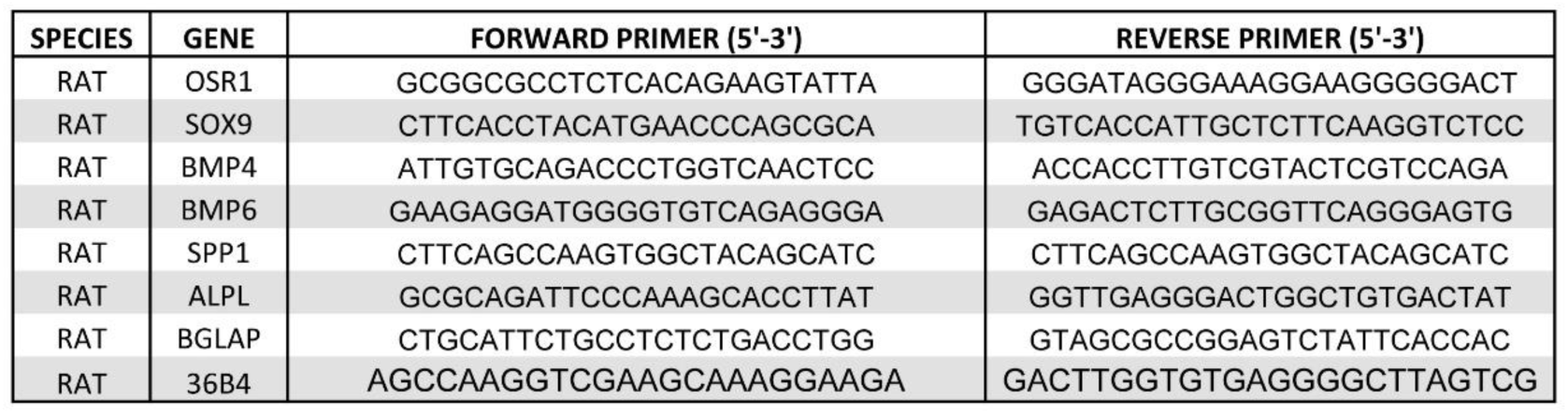
Table of qPCR primers.

**Figure 2:**
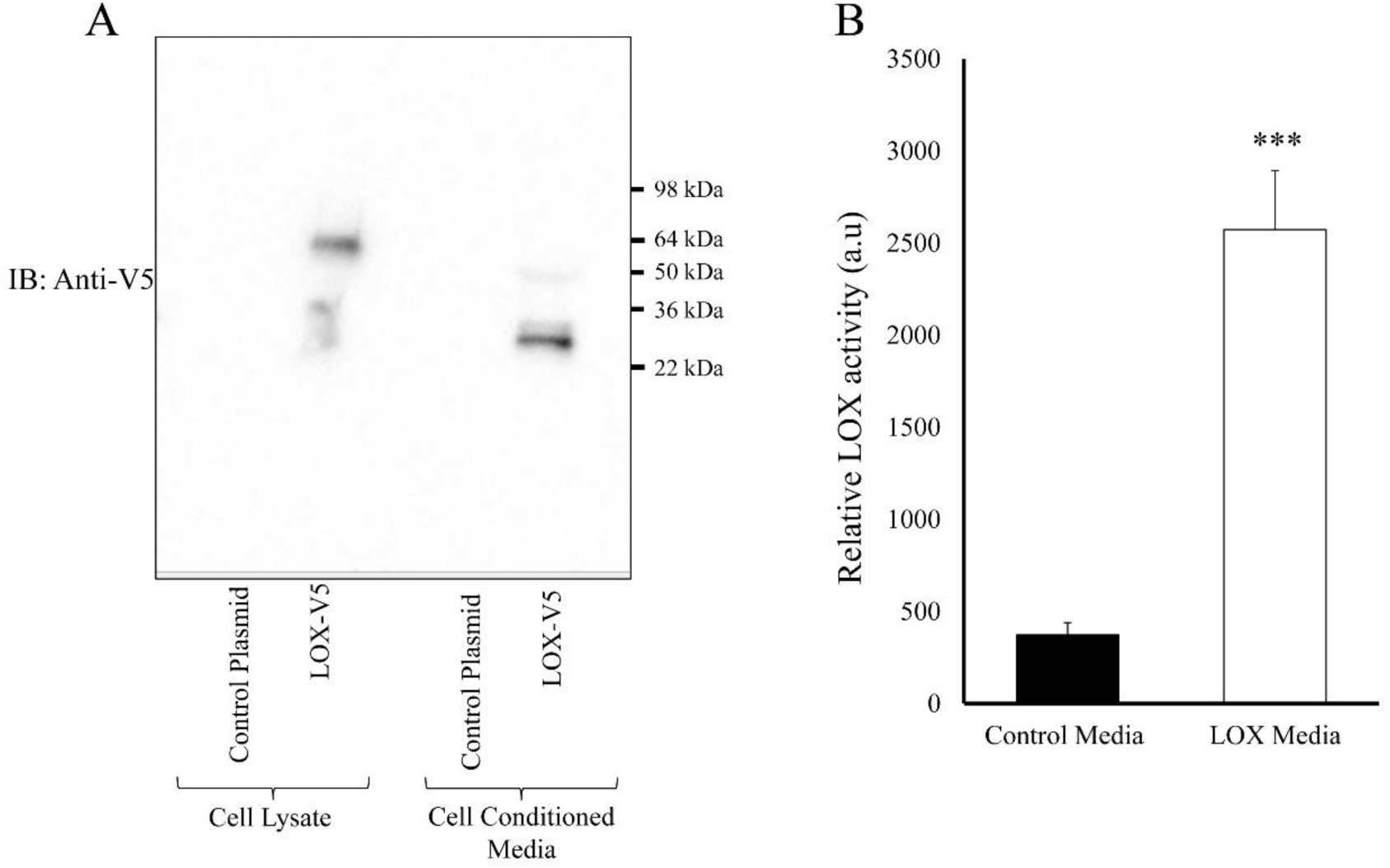
Validation of ecoptic LOX expression and activity. Rat VSMC were transfected with control plasmid or plasmid expressing LOX-V5. 24 hours post transfection, media was replaced with serum free media and conditioned for 6 hours. Ectopic LOX expression in total cell lysate of cell conditioned media was detected using western blotting with and anti-V% antibody (A). LOX activity was quantified in cell conditioned media (B). *** indicates p<0.001. One way ANOVA with Student Newman Keuls post-test.

**Figure 3:**
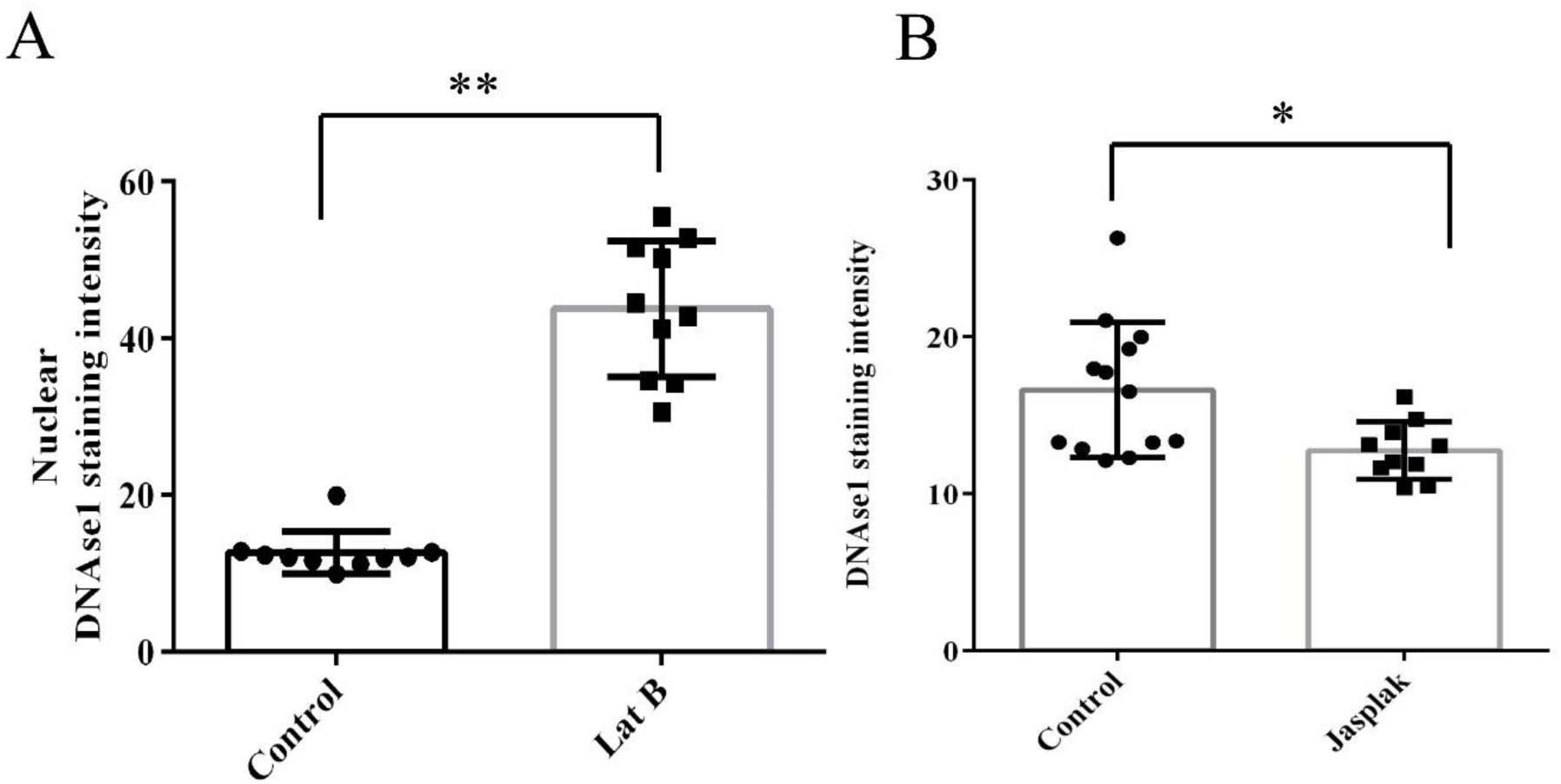
Validation of nuclear actin monomer detection using Alexa-Fluor 594-DNAse1 staining. Human saphenous vein samples were treated for 6 hours with 0.25 µg/ml latrunculin B (n=3; A) or 10 nM Jasplakinolide (n=3; B). Formalin fixed histological sections were stained with Alexa-fluor 594 conjugated DNAse1 to detect actin monomer. Nuclear levels of actin monomer were determined by confocal microscopy and image analysis using Image-J. * indicates P<0.05; ** indicates p<0.01. Students t-test

## Notes

### Competing Interest Statement

The authors have declared no competing interest.

